# From motif to function: Inferring the functions of long zinc finger proteins through combinatorial selection

**DOI:** 10.1101/2022.04.19.488842

**Authors:** Zheng Zuo

## Abstract

C2H2 zinc finger proteins (ZFPs) comprise of the largest group of DNA-binding proteins in human genome, and many of them contain long, tandem array of fingers, making the motif discovery, prediction of *in vivo cis*-regulatory elements (CREs), and understanding their functions particularly challenging. Previous work established that due to the dependent recognition between sub-motifs, the simple, additive recognition model impedes motif discovery and compromises our understanding about how ZFPs work. This work uses ZFP3, a 13-finger long ZFP with no known function, as case example to address the reverse question---given the full-length motif learned through *in vitro* experiments, like Spec-seq and HT-SELEX, how to reliably identify its *in vivo* cis-regulatory elements (CREs) and further predict this gene’s functions. Through sorting of all possible sites within the ChIP-seq peaks with similar predicted binding energy into groups and comparing the aggregate ChIP-seq signals between groups, it is evident that either its full-length or individual sub-motif alone fails to correctly identify all high-affinity specific sites without false-positives, thus it is necessary to revise current algorithm, and use both the core and upstream motifs as separate components to improve the prediction accuracy. Furthermore, significant number of regulatory elements of ZFP3 are found to be proximal to genes associated with microtubules organization and ciliogenesis, which coincides with the fact that ZFP3 is specifically upregulated in multiple ciliated cells. At last, local chromatin accessibility and active chromatin marks like H3K27ac are found to positively associate with the differential binding of ZFP3 between tested cell lines. Overall, this work establishes a novel “From motif to function” strategy for long ZFPs, and the data analysis workflows are implemented through R package TFCookbook for reuse onto other ZFPs.

## Introduction

As the largest group of DNA-binding proteins within human genome conferring diverse functions like genome insulation, recombination, and transposon silencing, C2H2-type ZFPs^1^ are unique in their long, tandem fingers configuration and therefore are associated with two unresolved problems.

First, among nearly 800 C2H2 ZFPs, around half of them contain KRAB domains and are perceived to be transcriptional repressors^2^. It is true that KRAB-containing ZFPs (KZFPs) carry more fingers per gene than those non-KRAB ZFPs on average, most likely driven by the evolutionary pressure to specifically silence those repetitive elements^3^, which emerged during the human evolutionary history, including LINEs, SINEs, LTRs, etc. One of my recent works has experimentally demonstrated that for some long KZFP like ZNF675, this exceedingly long-finger architecture is required to facilitate preferential recognition of certain LTR elements over non-repeat elements. However, there are still a lot of long, non-KRAB ZFPs, e.g., ZFX, ZFY, PRDM3, PRDM16, ZFP3, which are transcriptional activators and not involved in repeat silencing processes. It is unclear what is the driving force to maintain their long-finger architectures. Some recent work about ZFX^4^ claims that only small number of its fingers are sufficient to facilitate its biological functions. If so, why ZFX has 13 fingers and keeps evolutionarily conserved among mammals? Is it true that current partial motif information for most ZFPs^5^ are sufficient to understand their functions? More careful study is needed to test whether those seemingly “redundant” fingers contribute to the recognition of CREs and thus are essential for their functions.

Second, my previous work^6^ on ZFY and CTCF quantitatively demonstrated that the additive, position-independent recognition model^7^ fails to properly address the complex, dependent recognition relationship between sub-motifs within those long ZFPs. The blind use of position-weight matrix (PWM) should be cautioned. The lingering questions are how to include the dependency effect into current *in vivo* binding sites prediction algorithm and whether or not that inclusion can improve the accuracy of current algorithm.

ZFP3^8^, as a 13-finger long ZFP found in mammals, has no known function currently. Sequence alignment across eight mammals (Fig. 1A) reveals that its first ten fingers are highly conserved, whereas the N-terminal acidic-rich domain is divergent, which is common for other transcription activators like ZFX and ZFY. ENCODE consortium^9,10^ has generated ChIP-seq data of ZFP3 in HEK293 and SK-N-SH cell lines, each with reproducible replicates. In a separate large-scale motif discovery work based on existing ChIP-seq data, BPnet-based algorithm^11^ predicted some 28-nt long motif, which was later confirmed by Spec-seq experiment to large extent. With ChIP-seq, Spec-seq, and HT-SELEX data at hand, it serves as a very good study case to address two above problems and can demonstrate some general, effective strategy to predict the unknown functions of other ZFPs.

**Figure 1.**
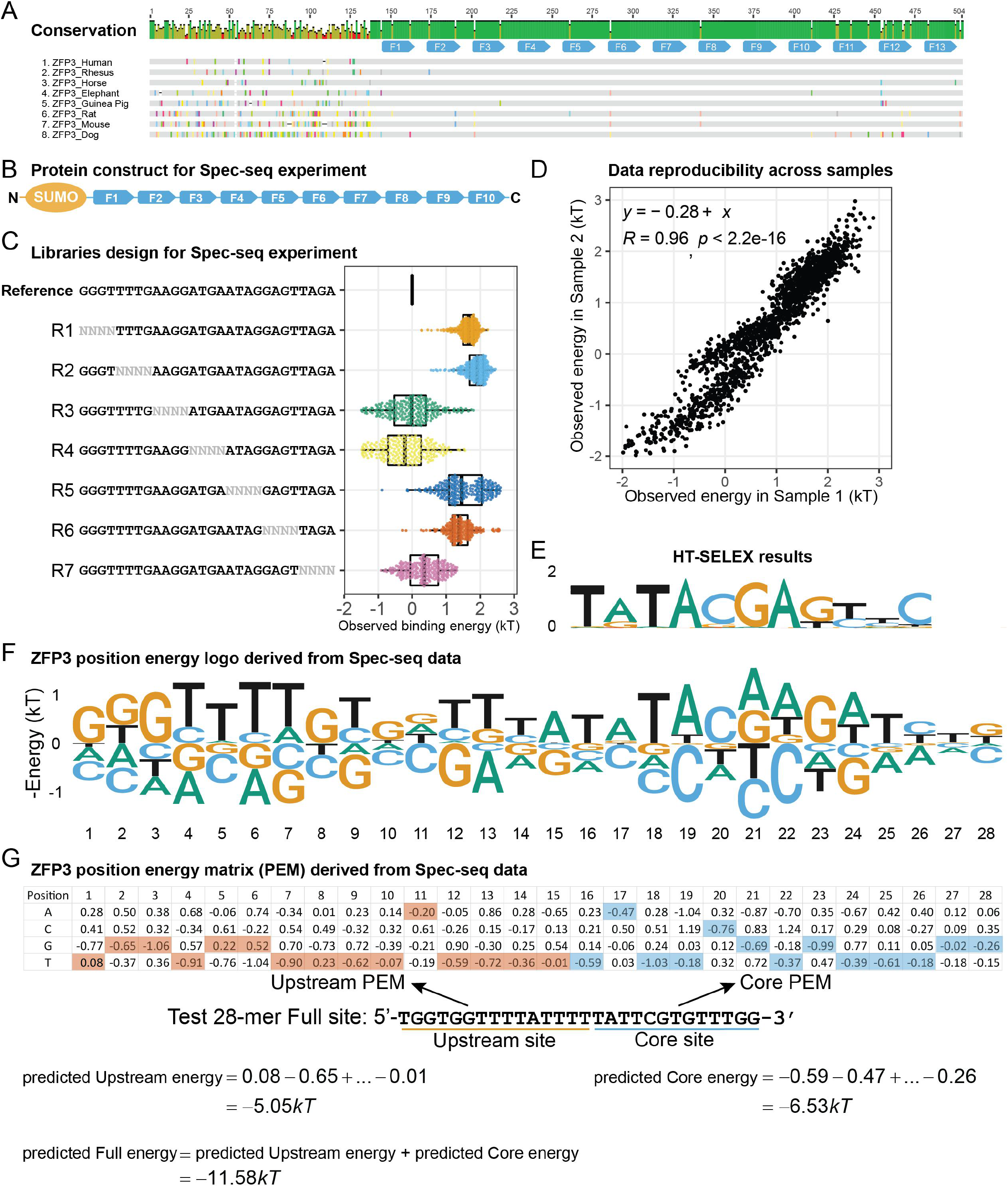
A) The sequence alignment of ZFP3 from eight mammals species; B) Protein construct used for ZFP3 Spec-seq experiment; C) The libraries design for Spec-seq experiment and the energy distribution of observed variants sorted by library type; D) Data reproducibility between replicates; E) The HT-SELEX derived motif for ZFP3; F) ZFP3 position energy logo derived from Spec-seq data (Sample 1), which corresponds to the PEM; G) Position energy matrix derived from Spec-seq data (Sample 1), which can be partitioned into two parts, i.e., the Upstream and Core motifs; The binding energy of some 28nt-long test site can be predicted additively.

## Results

### The full-length motif of ZFP3 can be derived from Spec-seq data reproducibly

Spec-seq^12^ is a medium throughput experimental method to quantitatively measure the specificity profile of some transcription factor (TF) of interest with resolution down to 0.2kT between replicates. Limited by sequencing capacity and some other factors, the number of tested binding sites in one assay cannot exceed a few thousands, therefore for long ZFP like ZFP3, some prior information about its putative consensus site is preferred for libraries design. Based on the 28nt long reference site inferred by BPnet algorithm, it becomes feasible to design a series of tandem, nonoverlapping degenerate libraries, namely R1 to R7, to systematically tile and perturb the whole reference site for Spec-seq work (Fig. 1C). Based on sequence conservation, the first ten fingers were included in the recombinant protein construct (Fig. 1B).

The default motif modeling method for Spec-seq data is essentially to find a set of parameters to minimize the total deviation between predicted and observed energy values for some chosen set of sequences, which has been implemented by R package TFCookbook^13^. Given that set of parameters in the form of position energy matrix (PEM), we can predict the binding energy of any other sequences using the protocol illustrated in Fig. 1G. It turned out the PEM and position energy logo derived from Spec-seq data are similar to the original BPnet prediction. Both the measured data and derived PEM parameters are highly concordant and reproducible between replicates (Fig. 1D, S1).

One thing slightly surprising is that the HT-SELEX derived motif is shorter and only matches the downstream part of fulllength PEM, which has been seen in other cases like CTCF and ZFY before^6^. If some sub-motif shows up in every single instance of specific binding site, that sub-motif is usually designated as core motif, while the flanking sub-motif is named as either upstream or downstream motif depending on its relative position to the core site^14^. For ZFP3, we can artificially divide the full-length PEM into two components, i.e., the upstream and core motifs (Fig. 1F). Using these two sub-motifs, we can predict the upstream site and core site energy correspondingly.

One implicit assumption about the usage of additive PEM is that some specific protein-DNA complex is already formed, and we only need take account the local perturbative effect of the base at each position. However, due to the complex dependency between sub-motifs reported in other ZFPs, the predicted energy value could be far from the real value, particularly for those sites containing defective core sequences. It is to be tested whether or not those predictions through simple, additive PEMs are fully consistent with *in vivo* ChIP-seq observations.

### Neither the full-length nor individual sub-motif can correctly predict all high-affinity, specific sites without false-positives

Since the ChIP-seq data are reported as sets of genomic peaks of length 200-700bp long and continuous spectrum of read-depth normalized signals across the whole genome, we have no prior knowledge how to attribute the observed ChIP-seq signals to individual specific binding sites within the peaks. Using individual sub-motif and full motif derived from Spec-seq data, it is feasible to predict the core, upstream, and full binding energy of every possible 28mer site within those reported peaks respectively and plot their overall distributions like Fig. 2A and 2E. Besides the binding site strength, many confounding factors, biologically and technically, can influence the observed ChIP-seq signals around individual binding site. To reliably inspect the consistency between predicted binding strength and observed ChIP-seq signals, we can sort those sites with similar predicted values into groups, align the ChIP-seq signals around each site (Fig. 2B, 2D), and compare the average, aggregate ChIP-seq signals between groups (Fig. 2C, 2F). Unless there is some other factor systematically biased toward a particular group and that factor is known to significantly affect ChIP-seq signals, the intensity of average, aggregate ChIP-seq signals should be proportional to the binding affinity of underlying sites within each group. Based on the analysis of available DNase-seq and various histone ChIP-seq signals, such bias factor is currently not found (Fig. S2). Note that even if we randomly pick one site from each ChIP-seq peak and calculate the aggregate ChIP-seq signals, we still observe some summit-like enrichment around the center (Fig. 2C, 2D, 2G), which certainly doesn’t mean this random selection is an effective way to identify the specific sites but tells us the randomly picked sites are probably proximal to some true specific sites within the peak. This random selection signal serves as negative control to tell us how much better any select group of sites are more effectively predicting the true specific elements.

**Figure 2.**
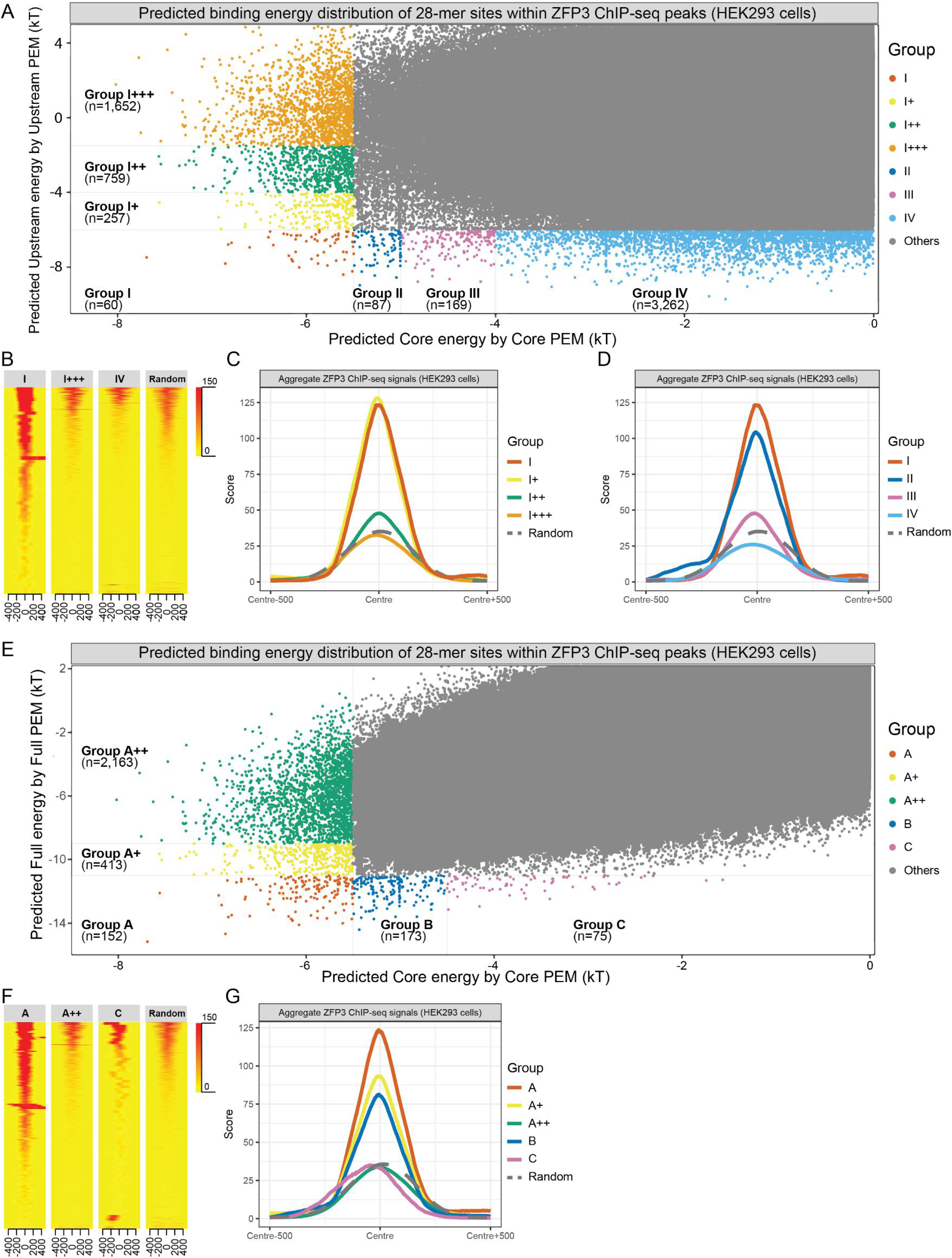
**A)** Distribution of all possible 28nt-long binding sites within ChIP-seq of HEK293 cells, sorted and classified by predicted Upstream and Core energy values; **B)** ChIP-seq signals aligned over binding sites within Group I, I+++, IV respectively; **C)** Average, aggregate ChIP-seq signals of Group I, I+, I++, and I+++; **D)** Average, aggregate ChIP-seq signals of Group I, II, III, and IV; Randomly selected sites serve as negative control. **E)** Distribution of all possible 28nt-long binding sites within ChIP-seq of HEK293 cells, sorted and classified by predicted Full and Core energy values; **F)** ChIP-seq signals aligned over sites within Group A, A++, and C; **G)** Average, aggregate ChIP-seq signals of Group A, A+, A++, B, C; Randomly selected sites serve as negative control.

We can sort all those 28mers within ChIP-seq peaks based on the predicted upstream and core energy values separately (Fig. 2A). It is possible to artificially divide some sites into Group I to IV based on their Core site strength (the lower energy, the higher affinity and binding strength), and divide some other sites into Group I to I+++ based on their upstream sites strength (Fig. 2A). Hypothetically, if only a small set of fingers recognizing the core motif are needed to facilitate proper recognition to the underlying targets and the upstream motif has no role, then there should be no ChIP-seq signals difference between Group I, I+, I++, and I+++, since they all have the same level of core sites strength. However, we do see incrementally decreasing signals from Group I to Group I+++(Fig. 2C), proving that the upstream motif does contribute to the specific recognition of underlying targets. To consistently estimate the prediction accuracy of various selection methods, we can classify all those groups of sites with at least 50% of highest group signal as high-affinity, specific sites and other groups as low-affinity sites. For the case of HEK293 data, only Group I, I+, and II are high-affinity sites. If we use the core motif alone as the selection criteria *E_Core_* <– 5*kT* to pick high-affinity sites, then Group II sites are excluded from that selection, while Group I+++ sites are included, though Group I+++ sites elicit no stronger ChIP-seq signals than the negative control of randomly selected sites. The sensitivity of this Core-motif-only selection method is no higher than

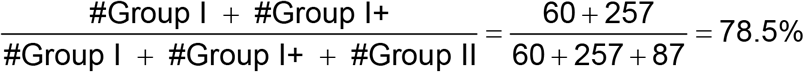

and the specificity (here defined to be the percentage of true specific sites in prediction) is only

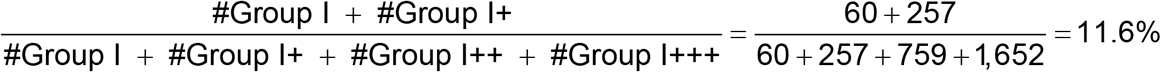

Similarly, if we perform Upstream-motif-only selection based on criteria *E_Upstream_* <– 6*kT*, Group I+ sites are excluded even though their ChIP-seq signals are significantly stronger than Group IV signals. In fact, the Group IV sites signals are even weaker than the random selection control, meaning that this group is predominantly composed of non-specific sites. The sensitivity and specificity of this selection are estimated to be no better than 36.4% and 4.1% respectively.

Alternatively, we can sort all those 28mers within ChIP-seq peaks based on their predicted full and core energy values (Fig. 2E), it is possible to manually divide some sites with the same full energy levels into Group A to C based on their core site strength, and divide some other sites with the same core energy levels into Group A to A++ based on their Full site strength (Fig. 2E). Theoretically, if simple, additive recognition model suffices to predict the binding site strength of all observed sites, group A, B and C should exhibit the same level of ChIP-seq signals, since they are all predicted to have the same full-site strength. However, Group C sites elicit no stronger signals than random selection, most likely because they contain defective core sequences. This observation is consistent with the conclusion of previous work about the dependent recognition of long ZFPs, i.e., the proper recognition of flanking sequence depends on the presence of intact core site, otherwise the whole site is in non-specific state regardless of the degree of flanking sequence match. With the same rules, Group A, B, A+ are classified as high-affinity sites, and others are low-affinity sites. Under the Full-motif-only selection criteria *E_Full_* <– 11*kT*, the detection sensitivity and specificity cannot exceed 44% and 81.3% respectively. It is true we could increase the detection specificity by further lowering the selection threshold to −12 kT, but in that case, many more high-affinity sites within Group A are excluded and the detection sensitivity would drop even lower. These results are based on ChIP-seq data in HEK293 cells, and if we do the same analysis on SK-N-SH cells, the same conclusion can be drawn (Table 1, Fig. S3).

**Table 1.** Estimated sensitivity and specificity of various selection methods for high-affinity, specific sites

| Selection method | HEK293 cells |  | SK-N-SH cells |  |
| --- | --- | --- | --- | --- |
|  | Sensitivity | Specificity | Sensitivity | Specificity |
| Core-motif-only selection ($E_{Core} < -5kT$) | 78.5% | 11.6% | 86.2% | 6.2% |
| Upstream-motif-only selection ($E_{Upstream} < -6kT$) | 36.4% | 4.1% | 24.7% | 1.5% |
| Full-motif-only selection ($E_{Full} < -11kT$) | 44% | 81.3% | 31.6% | 70.1% |

### Combinatorial selection of high-affinity sites with separate motif components excludes spurious, non-specific sites

Based on above observations, we can conclude that full motif or individual sub-motif alone fails to correctly select for all high-affinity sites without false positives like group C, group III, or group I+++. To improve the identification accuracy and include the top three group of sites into selection, we need use at least two separate components like the predicted upstream and core sites energy to do combinatorial selection, e.g., *E_Upstream_* <– 4*kT*, *E_core_* < −5*kT*. Clearly under this criterion, those weakly specific sites in Group I+++ and spurious, non-specific sites in Group C are excluded, so both the sensitivity and specificity are certainly superior to three other mentioned methods.

**Table 2.**
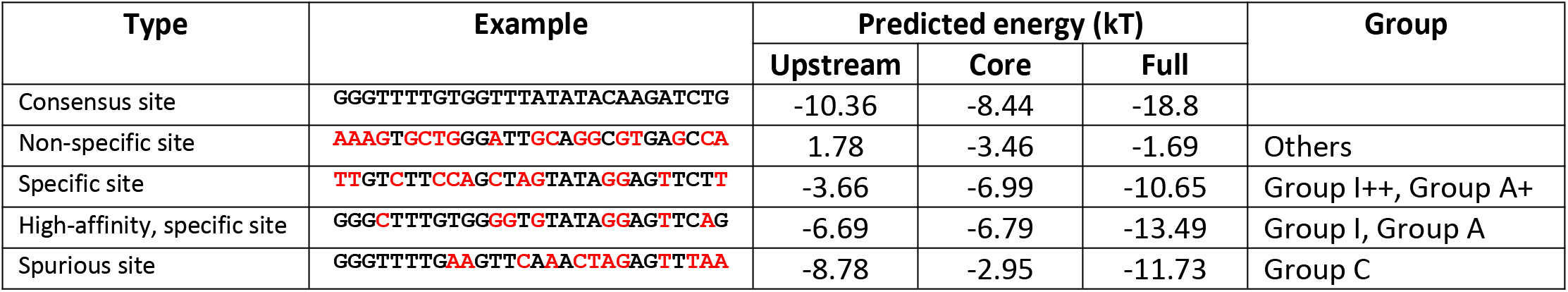
Examples of different types of binding sites

### Significant number of high-affinity, specific sites are associated with genes of microtubule organization and ciliogenesis

Current ChIP-seq data is acquired under the condition of ZFP3 overexpression, but the endogenous expression levels of ZFP3 are much lower (Fig. 3D). Under the endogenous condition, this TF is expected to be more localized to those high-affinity sites and other sites are depleted, rather than distributed broadly, so we can reasonably assume its functional CREs are largely within high-affinity, specific sites (Fig. 3B). Among total 2,154 high-affinity sites identified in HEK293 and SK-N-SH cell lines, 111 sites are found to be proximal to the transcription start sites (T.S.S.) of some genes with no more than 500bp distance, namely T.S.S. proximal elements. Through annotation of associated genes with ChIPpeakAnno^15^, it is evident that the top three hits (FSD1, MARK1, and SPC24) are directly involved in the microtubule organizations. Interestingly, many other hits, including CEP95, STOML3, TTC26, CEP162, CERKL, and MAP11, are involved in the ciliogenesis process, which intimately depends on the proper microtubule organization. No other genes with specific functions are found enriched. According to the single-cell gene expression data compiled by the Human Protein Atlas^16^, ZFP3 is specifically upregulated in multiple ciliated cells from diverse tissue types, including lung, bronchus, eye, and testis. It is highly unlikely that these two observations are pure coincidences, thus ZFP3 is inferred to be the positive transcription regulator of microtubule organization and ciliogenesis processes.

**Figure 3.**
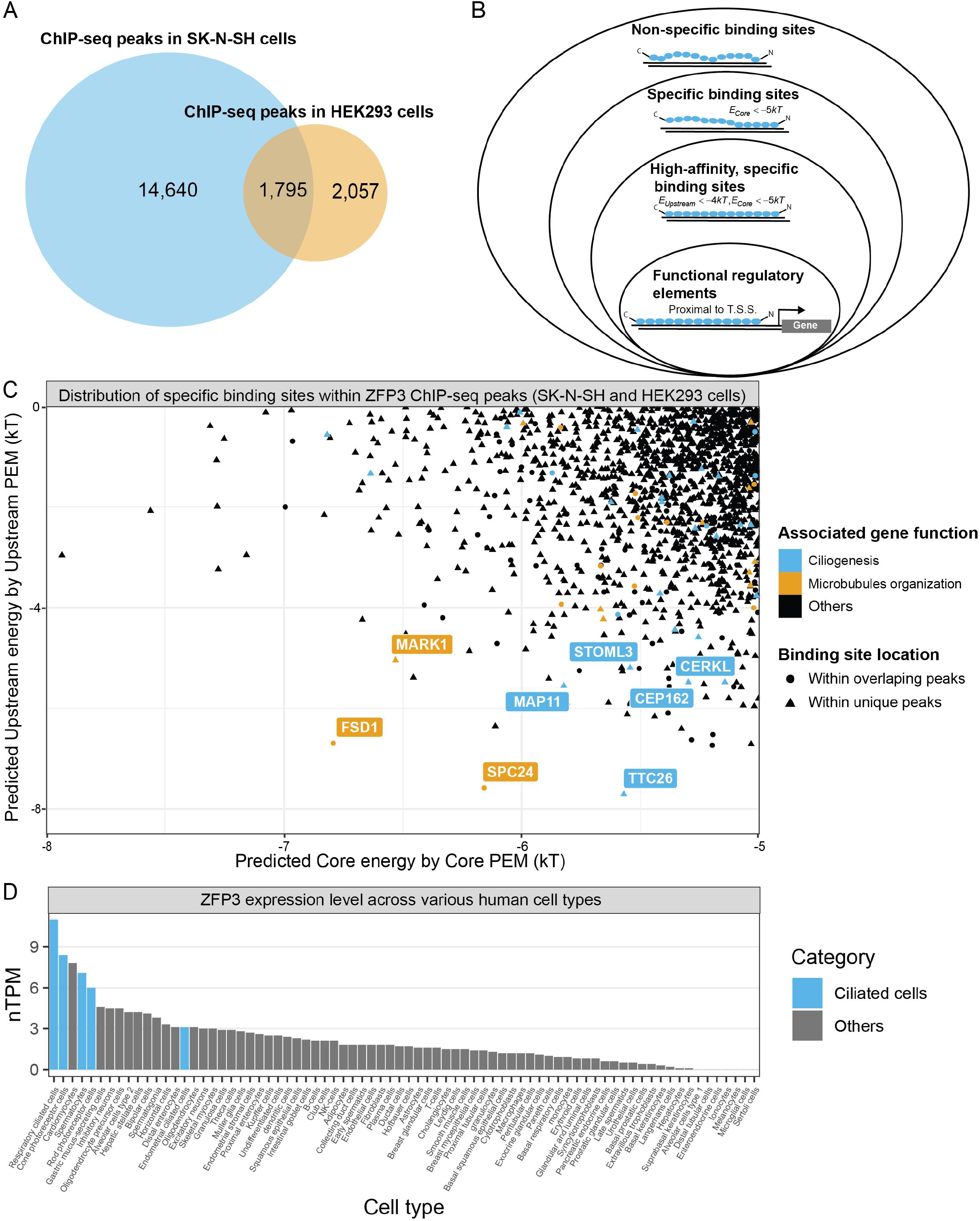
**A)** The Venn diagram illustration for the overlap between ChIP-seq peaks of ZFP3 in HEK293 and SK-N-SH cells; **B)** The classification and search criteria of functional CREs within high-affinity, specific sites; **C)** The distribution of specific sites found within both cell types; Only T.S.S. proximal elements (no more than 500nt) are shown; **D)** ZFP3 expression level across various human cell types (Source: Human Protein Atlas).

Most of currently identified ZFP3 targets are structural components of cytoskeleton or cilia structures. For example, FSD1 and CEP162 are essential for transition zone assembly^17,18^; SPC24 belongs to NDC80 kinetochore complex^19^; WDR19(IFT144) belongs to the IFT-A complex^20^; TTC26 belongs to IFT-B complex^21^; MAP11 belongs to TRAPP (transport protein particle) II complex^22^. Some target, SPG11, was shown to associate with microtubules and its dysfunction impairs cytoskeleton stability even without definitive structural annotation^23^. Three genes, MARK1, KAT2B and MOB3C, are of special interest, because they are known cytoskeleton regulators and the identified CREs of the first two are highly conserved according to PhastCon sequence alignment of 30 mammals (27 primates) (Fig. 5A). Recently, KAT2B was found to negatively regulate the centrosome amplification through its acetylase activity^24^. The maintenance of cytoskeleton structure and ciliogenesis are highly complex and certainly require delicate control and coordination on multiple levels.

Visual inspection of ChIP-seq signals reveals that many identified, specific elements are very close to the ChIP signal summits (Fig. 4, 5A), which lends further evidence to the accuracy of the revised algorithm. There is no one-to-one correspondence between peak and specific binding site, e.g., multiple specific sites have been identified near the MAP1S peak region (Fig. 4E). This kind of composite peak might elicit different induction effect of downstream gene under increasing ZFP3 concentration, which is worth further investigation.

**Figure 4.**
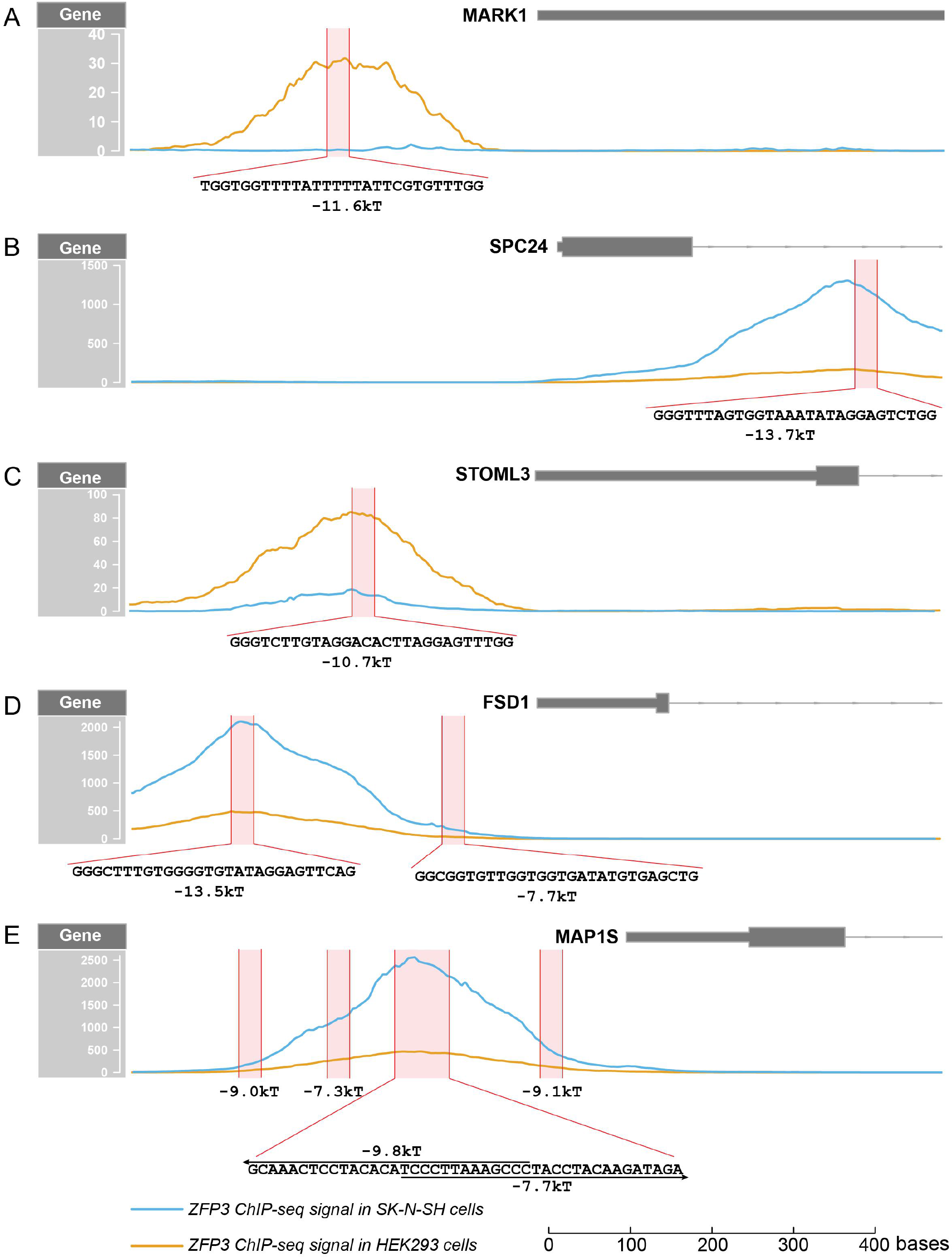
**A-E)** The ChIP-seq signals profiles for five identified regulatory targets. Specific sites with full-binding energy below −7kT are annotated as red boxes.

**Figure 5.**
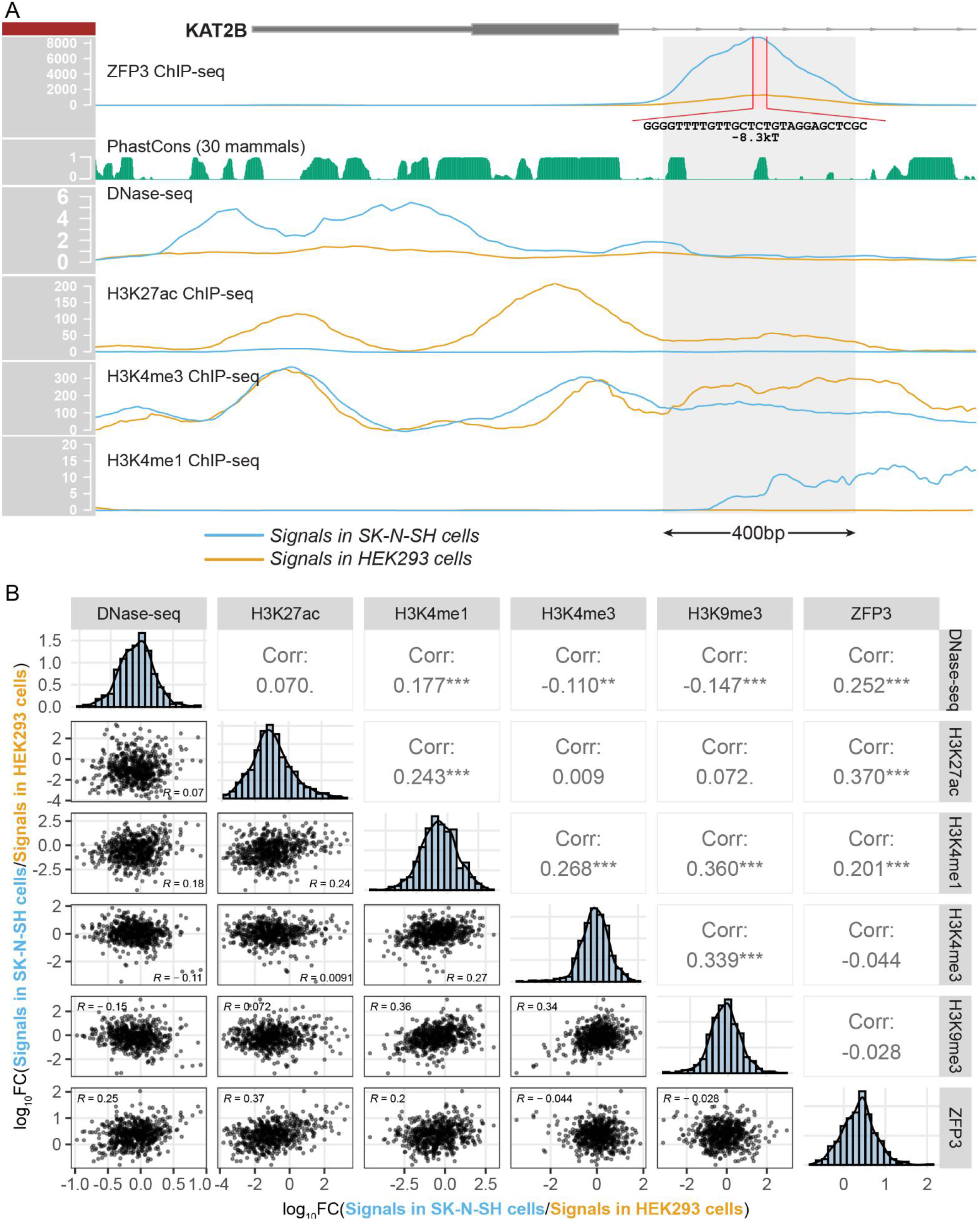
A) ChIP-seq, DNase-seq, and PhastCon signals near KAT2B promoter region; 400bp window around the identified specific binding site is annotated as shadows; B) Pairwise correlation analysis between the logFC of signals between SK-N-SH and HEK93 cells within the 400bp windows around 615 overlapping, high-affinity sites. ** indicates p-value < 0.01; *** indicates p-value < 0.001.

### Local DNase-seq and H3K27ac signals positively associate with the differential occupancy levels of ZFP3 between tested cell lines

Though the same experiment protocols and data processing pipelines were used for the ZFP3 ChIP-seq work, most of observed peaks don’t overlap between HEK293 and SK-N-SH cell lines (Fig. 3A). Even among ~1,800 overlapping peaks, the signal intensity can be very different from case to case (Fig. 4, 5A), meaning that binding site affinity is only necessary but not sufficient to yield strong ChIP-seq peak profiles. To quantitatively characterize the contributions of other potential confounding factors, we can use identified 615 overlapping, high affinity sites as reference elements, and quantify the fold change of signals of ZFP3 ChIP-seq, DNase-seq, and all available histone ChIP-seq data within some 400bp window around each reference element between two cell lines (Fig. 5). Pairwise correlation analysis between these sets of continuous variables reveals that DNase-seq, H3K27ac, and H3K4me1 positively associate with the preferential binding of ZFP3 between the two cell lines (R=0.25, 0.37, and 0.20 each with p-value < 0.001), while H3K4me3 and repressive chromatin mark H3K9me1 exhibit no significant correlations with ZFP3 ChIP-seq signals (Fig. 5B). Chromatin accessibility only partly explains the observed ChIP-seq signals difference. In fact, most of specific sites found within ChIP-seq peaks don’t overlap with the DNase-seq peaks of HEK293 and SK-N-SH cells (Fig. 5A, Fig. S4). We cannot conclude that active chromatin marks like H3K27ac directly recruit ZFP3 to nearby region, but it is very likely that those active histone marks are labels for favorable environments for ZFP3 binding. As more histone ChIP-seq signals become available on those reference cell lines, we should be able to do same kind analysis and find more contributing factors to the occupancy of long ZFPs with acidic activation domain like ZFP3.

## Discussion

This work is part of series of work about the form, mode, function, and diseases of long ZFPs and aims to resolve two problems mentioned in the Introduction section. ZFP3 is good example because we happen to have its full-length motif but have yet confirmed its biological functions.

For problem one, if we only know ZFP3’s motif partially like the core part derived from HT-SELEX results, which is common for many other long ZFPs, we wouldn’t be able to distinguish those high-affinity, specific binding sites from other specific sites only carrying good core sequences and further infer this gene’s functions. Besides repeat elements silencing, specifically activating certain functional regulatory elements provides another mechanism for ZFPs to keep such long-finger architecture. In short, those long ZFPs need to be long for alternative reason. It motivates us to characterize the specificity profile of those long ZFPs in more detail.

Second, as one anonymous reviewer on my previous work wisely pointed out, the value of dependent recognition model depends on whether or not it advances the knowledge of this field. Indeed, all models are wrong, some are useful (George Box’s words). If we stick to the use of simple, additive PWM to predict the *in vivo* CREs, no matter how long or accurate the motif is and how we adjust the selection threshold, we will always predict a lot of spurious, non-specific sites, like the Group C sites (Table 2, Fig. 2E), but paradoxically exclude many apparently high-affinity sites. To rationally exclude those spurious, non-specific sites, we must acknowledge that proper recognition to the core site is the priori for the formation of specific protein-DNA complex. In other words, dependent recognition model mechanistically gives us clue how to pick those high-affinity, specific sites. For ZFP3, combinatorial use of core and flanking motifs as separate components is the practical way to do that pick. This should be generally applicable to other long ZFPs.

In current work, all ZFP3’s CREs are searched within available ChIP-seq peaks regions of tested cell lines, but not those primary, ciliated cells. We may use open chromatin regions defined by DNase-seq or ATAC-seq signals in ciliated cells as the starting search regions to predict more targets, but current correlation analysis shows chromatin accessibility only plays a limited role in determining where ZFP3 can bind and regulate (Fig. 5, S4). The ultimate proof to confirm ZFP3’s functions comes from phenotypic evidence, either experimentally or clinically. Current literature and clinical genetics databases haven’t reported any severe diseases or deleterious phenotype associated with its loss of function (LoF), even though it is highly conserved across placental mammals. Since it is upregulated during the spermatogenesis process, it is likely that its LoF mutant would directly cause infertility and produce no viable offspring. To experimentally verify its functions, some organoid based system involving ciliogenesis process might be desirable.

## Acknowledgement

The ChIP-seq data of ZFP3 was generated at Michael Snyder lab and made available through ENCODE consortium; The initial full-length motif was inferred by BPnet-based algorithm developed by Anshul Kundaje lab; The Spec-seq data was produced by me at Polly Fordyce lab; The HT-SELEX result was produced by Tim Hughes lab at University of Toronto. All above results have been reported and submitted in separate paper. The ZFP3 expression data across cell types is accessed from Human Protein Atlas. This work was done after my departure from Stanford. I thank the helpful comment of anonymous reviewer in previous paper about the need to justify the value of dependent recognition model of ZFPs.

## Data availability and analysis workflow

The Spec-seq data for ZFP3 have been deposited at NCBI GEO database (#GSE189817). All ChIP-seq data are preprocessed by ENCODE consortium and are listed in Supplemental Key Resource table. The data analysis workflow for each figure have been deposited at GitHub (https://github.com/zeropin/ZFPCookbook/tree/master/ZFP3) for inspection and reuse. Identified specific elements of ZFP3 are listed in Supplemental Table.

