## Supplemental Figures for "From motif to function: Inferring the functions of long zinc finger proteins through combinatorial selection"

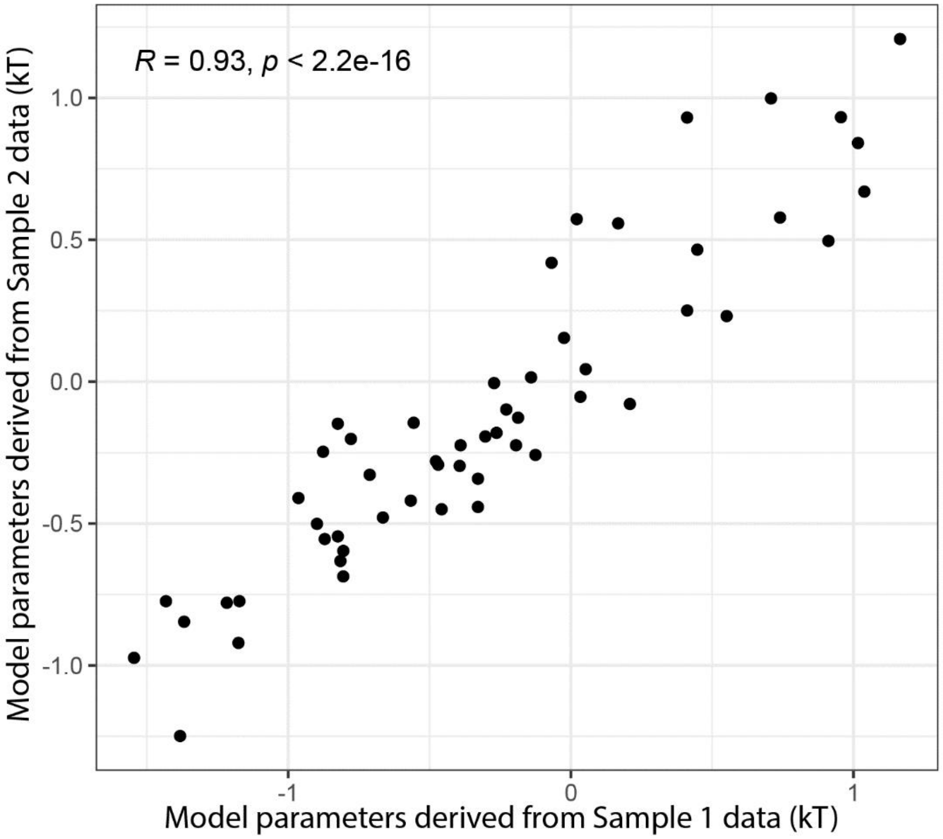


**Figure S1** Comparison of model parameters derived from replicate Spec-seq samples

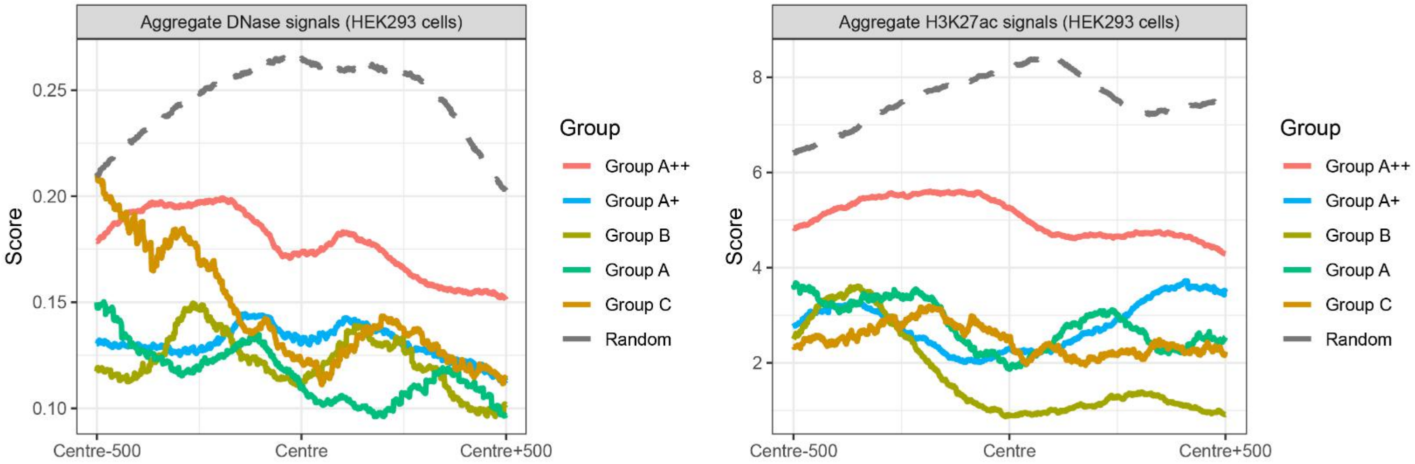


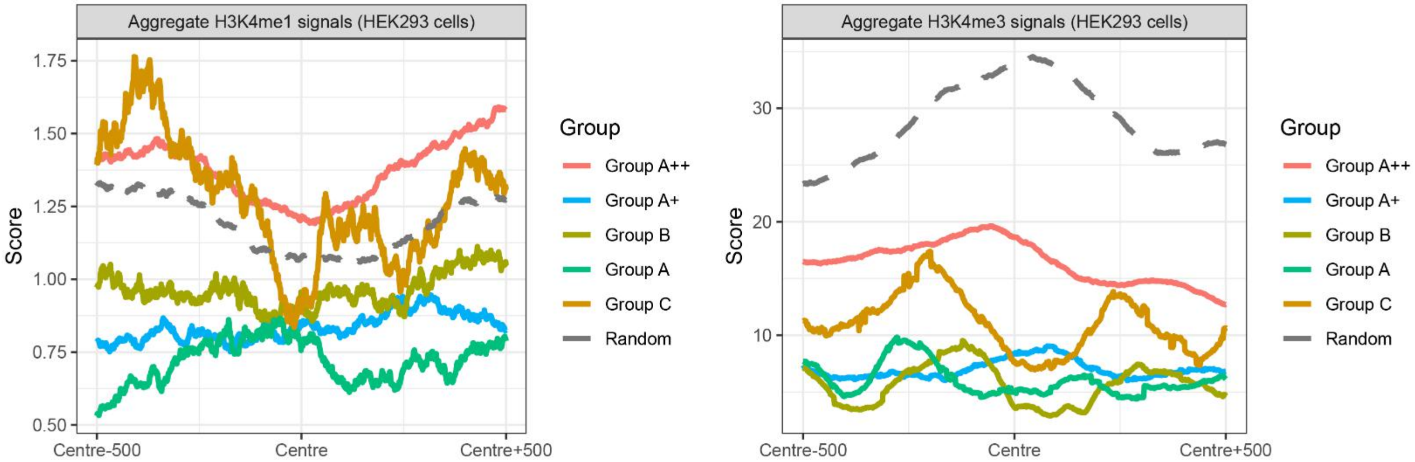


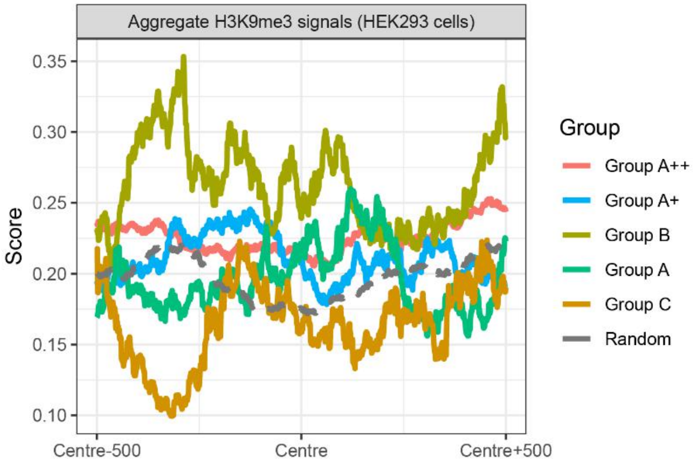


**Figure S2**. Aggregate signals for other potential coufounding factors sorted by Groups in Fig. 2E.
**A)** DNase-seq signals;
**B)** H3K27ac signals;
**C)** H3K4me1 signals;
**D)** H3K4me3 signals;
**E)** H3K9me3 signals;

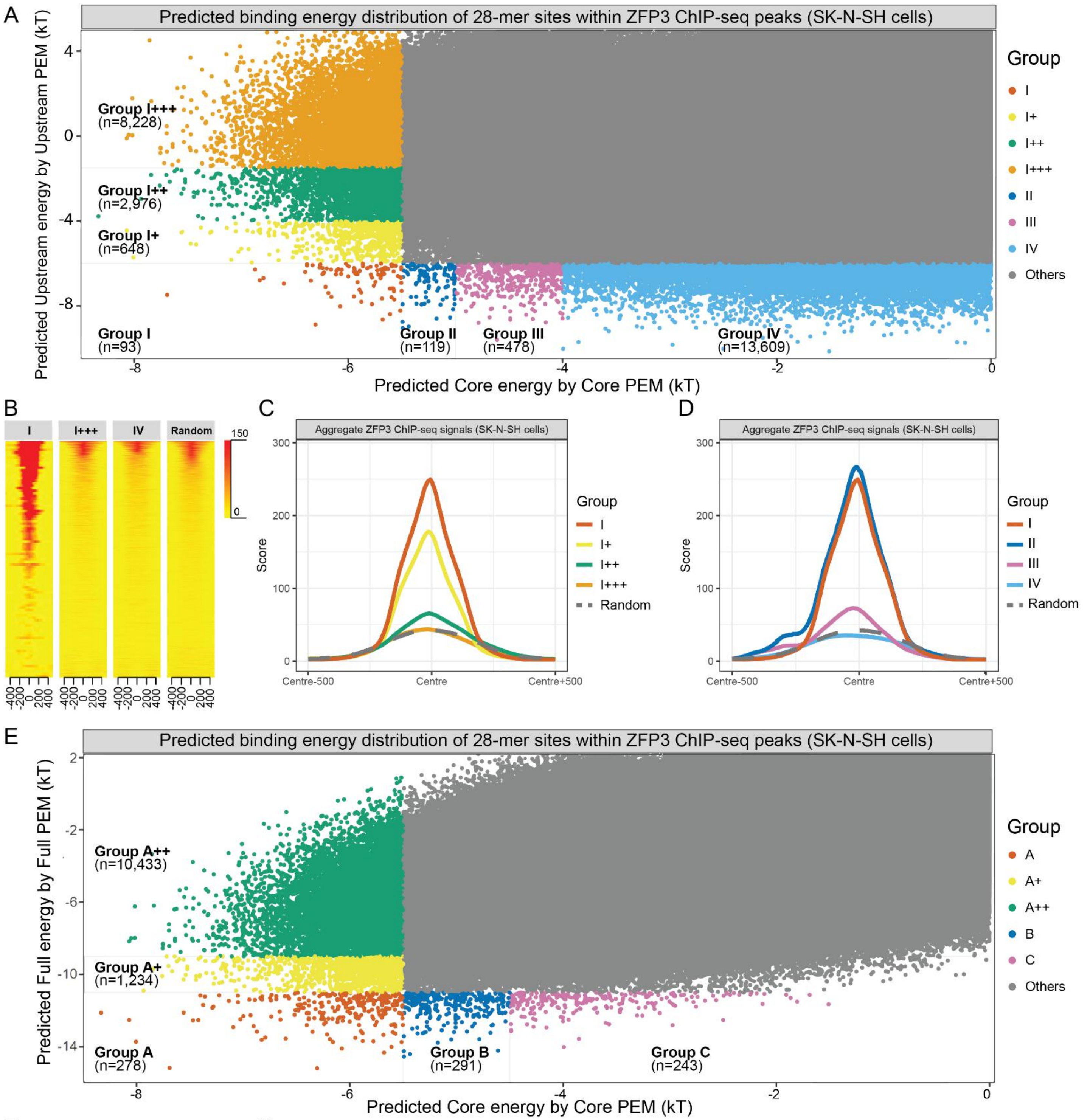


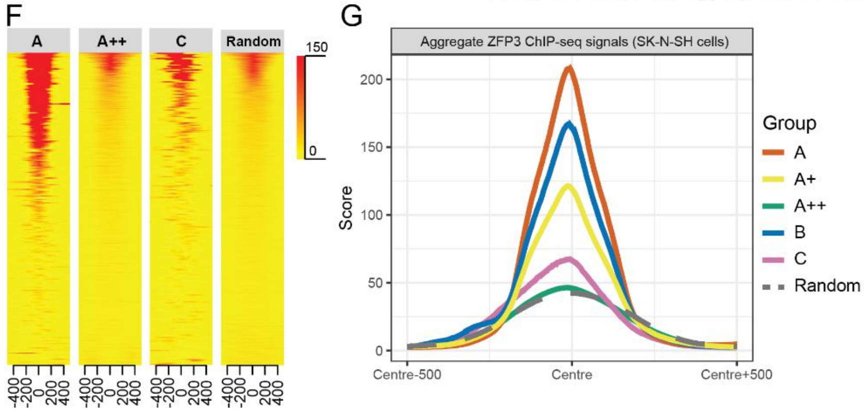


**Figure S3**

**A)** Distribution of all possible 28nt-long binding sites within ChIP-seq of SK-N-SH cells, sorted and classified by predicted Upstream and Core energy values;
**B)** ChIP-seq signals aligned over binding sites within Group I, I+++, IV respectively;
**C)** Average, aggregate ChIP-seq signals of Group I, I+, I++, and I+++;
**D)** Average, aggregate ChIP-seq signals of Group I, II, III, and IV; Randomly selected sites serve as negative control.
**E)** Distribution of all possible 28nt-long binding sites within ChIP-seq of SK-N-SH cells, sorted and classified by predicted Full and Core energy values;
**F)** ChIP-seq signals aligned over sites within Group A, A++, and C;
**G)** Average, aggregate ChIP-seq signals of Group A, A+, A++, B, C; Randomly selected sites serve as negative control.

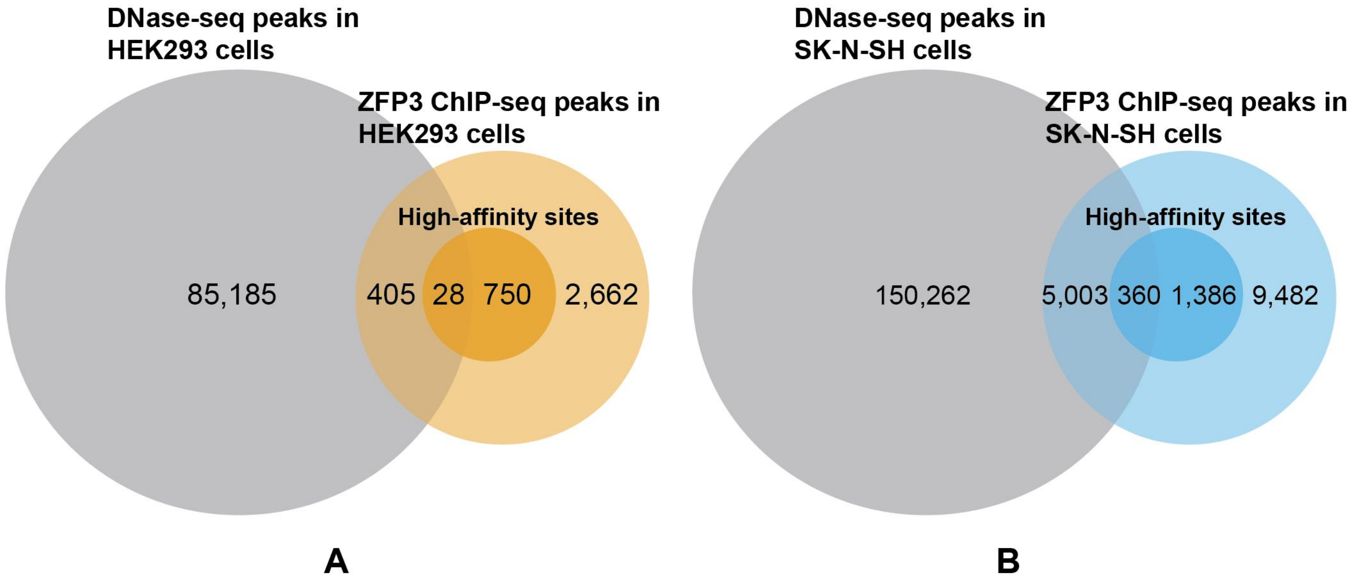


**Fig. S4**

**A)** Venn diagram illustration of the overlaps between DNase-seq peaks, ZFP3 ChIP-seq peaks, and high-affinity sites found within ChIP-seq peaks of HEK293 cells;

**B)** Venn diagram illustration of the overlaps between DNase-seq peaks, ZFP3 ChIP-seq peaks, and high-affinity sites found within ChIP-seq peaks of SK-N-SH cells.
